# New interfaces on MiD51 for Drp1 recruitment and regulation

**DOI:** 10.1101/324616

**Authors:** Jun Ma, Yujia Zhai, Ming Chen, Kai Zhang, Quan Chen, Xiaoyun Pang, Fei Sun

**Affiliations:** National Laboratory of Biomacromolecules, CAS Center for Excellence in Biomacromolecules, Institute of Biophysics, Chinese Academy of Sciences, 15 Datun Road, Beijing 100101, China.; State Key Laboratory of Biomembrane and Membrane Biotechnology, Institute of Zoology, Chinese Academy of Sciences, Beijing 100101, China.; University of Chinese Academy of Sciences, 19 Yuquan Road, Beijing 100049, China.

**Keywords:** Mitochondrial fission, Drp1, MiD51, dimerization, regulation

## Abstract

Mitochondrial fission is facilitated by dynamin-related protein Drp1 and a variety of its receptors. However, the molecular mechanism of how Drp1 is recruited to the mitochondrial surface by receptors MiD49 and MiD51 remains elusive. Here, we showed that the interaction between Drp1 and MiD51 is regulated by GTP binding and depends on the polymerization of Drp1. We identified two regions on MiD51 that directly bind to Drp1, and found that dimerization of MiD51 via an intermolecular disulfide bond between C452 residues is required for MiD51 to directly interact with Drp1. Our Results have suggested a multi-faceted regulatory mechanism for the interaction between Drp1 and MiD51 that illustrates the potentially complicated and tight regulation of mitochondrial fission.

## INTRODUCTION

Mitochondria are highly dynamic organelles that constantly undergo fusion, fission and move along the cytoskeleton ^1^. Beyond the primary function of mitochondrial dynamics in controlling organelle shape, size, number and distribution, it is clear that dynamics are also crucial to specific physiological functions, such as cell cycle progression, quality control and apoptosis ^2–5^. Dysfunction in mitochondrial dynamics has been implicated a variety of human diseases, including neurodegenerative diseases, the metabolism disorder diabetes and cardiovascular diseases ^6,7^.

Mitochondrial fission is mediated by multi-factors, such as dynamin-related protein Drp1 (Dnm1p in yeast) and its receptors on mitochondrial outer membrane, dynamin-2 (Dyn2) and endoplasmic reticulum ^8, 9^. However, Drp1 protein is mostly localized in the cytoplasm and must be recruited to the mitochondria by receptors on the mitochondrial outer membrane in response to specific cellular cues ^10^. After targeting, Drp1 self-assembles into large spirals in a GTP-dependent manner and then contributes to mitochondrial membrane fission via GTP hydrolysis ^5, 11^. In yeast, the integral outer membrane protein fission protein 1 (Fis1) interacts with two adaptor proteins, Caf4 and Mdv1, providing an anchoring site for Dnm1p recruitment. In mammals, three integral outer membrane proteins, Mff, MiD51 and MiD49, were identified as receptors recruiting Drp1 to mitochondria. Overexpression of Mff induces Drp1 recruitment and mitochondrial fission ^12–14^. MiD51 and MiD49 are anchored in the mitochondrial outer membrane via their N-terminal ends, and most of the protein is exposed to the cytosol. MiD51 and MiD49 specifically interact with and recruit Drp1 to mitochondria and then facilitate Drp1-directed mitochondrial fission ^15^. It is notable that the expression of both MiD49 and MiD51 appears to be up-regulated in pulmonary arterial hypertension (PAH), one characteristic of which is rapid cell division associated with Drp1-mediated mitochondrial division ^16^. And knock-down of endogenously elevated levels of MiD49 or MiD51 induces mitochondrial fusion ^16^.

Crystal structures of the cytosolic domains of MiD49 and MiD51 were reported and indicate that these proteins possess nucleotidyltransferase (NTase) folds and belong to the NTase family ^17, 18^. However, both proteins lack the catalytic residues required for transferase activity ^17, 18^. MiD51 does bind adenosine diphosphate (ADP) as a cofactor, but MiD49 lacks this capacity. The recruitment of Drp1 to the mitochondrial outer membrane by MiD51 was also addressed by two studies ^17, 18^ where a single exposed loop corresponding to residues 238-242 on the surface of MiD51 was identified as the Drp1-binding loop. Mutants lacking this active loop are defective in recruiting Drp1 to the mitochondrial surface. But there are still paradoxical and unclear aspects about the molecular mechanisms of Drp1 recruitment^17, 18^. In addition, Lóson *et al* ^18^proposed that MiD51 forms a dimer mainly via electrostatic interactions within the N-terminal segments and that dimerization is required for MiD51 mitochondrial fission activity but not Drp1 recruitment. Dimerization of MiD51 was not even addressed in Richter *et al*’s work^17^. Moreover, it is still not clear how the fission activity of MiD51 is co-regulated with Drp1.

Here, by combining structural biology, biochemical and biophysical techniques, we reveal that the interaction between MiD51 and Drp1 is regulated by the nucleotide acid binding state and polymerization of Drp1, and identify a second region on MiD51 that is important for Drp1 binding. We also show that MiD51 can form a homodimer through intermolecular disulfide bonds between the C452 residues, and it’s essential for direct interaction of MiD51 with Drp1. These results provide further insight into the molecular mechanism of interaction between Drp1 and MiD51, which plays key roles in mitochondrial fission regulation.

## RESULTS

### Interaction between MiD51 and Drp1 is dependent on the oligomerization of Drp1

It was reported that the N-terminus of MiD51 is anchored in the mitochondrial outer membrane and the C-terminus is cytosolic (Fig. S1A). The cytosolic domain MiD51^133-463^ can fold into a compact structure and are required for binding to Drp1^17, 18^. Considering that MiD51 does not undergo a conformational change upon ADP binding and mutants defective in ADP binding are still capable of recruiting Drp1 ^18^, we speculate that the roles of ADP or its analogues in regulating MiD51 are independent of Drp1 binding. But Drp1 is a GTPase protein belonging to the Dynamin super-family, and it can bind and hydrolyze GTP. We speculated whether MiD51 could selectively and dynamically recruit Drp1 under different nucleotide states. To test this, we performed GST pull-down assays in the presence of GTP, GDP, GDP+AlF_X_ and non-hydrolyzed GTP analogs GMP-PNP. We found that MiD51^133-463^ interacts with the GMP-PNP-bound state of Drp1 with high affinity, and this affinity is even higher than for the GTP bound state (Fig. 1A). Under GDP+AlF_X_ and GDP conditions, the strength of the interaction between MiD51^133-463^ and Drp1 is almost the same as for the apo state (Fig. 1A). However, the K38A mutant of Drp1, which is defective in nucleotide hydrolysis ^19^, appears to have no difference in binding affinity to MiD51^133-463^ under different nucleotide states (Fig. 1A). Based on these results we conclude that the interaction between MiD51 and Drp1 undergoes changes during the process of GTP hydrolysis.

**Figure 1.**
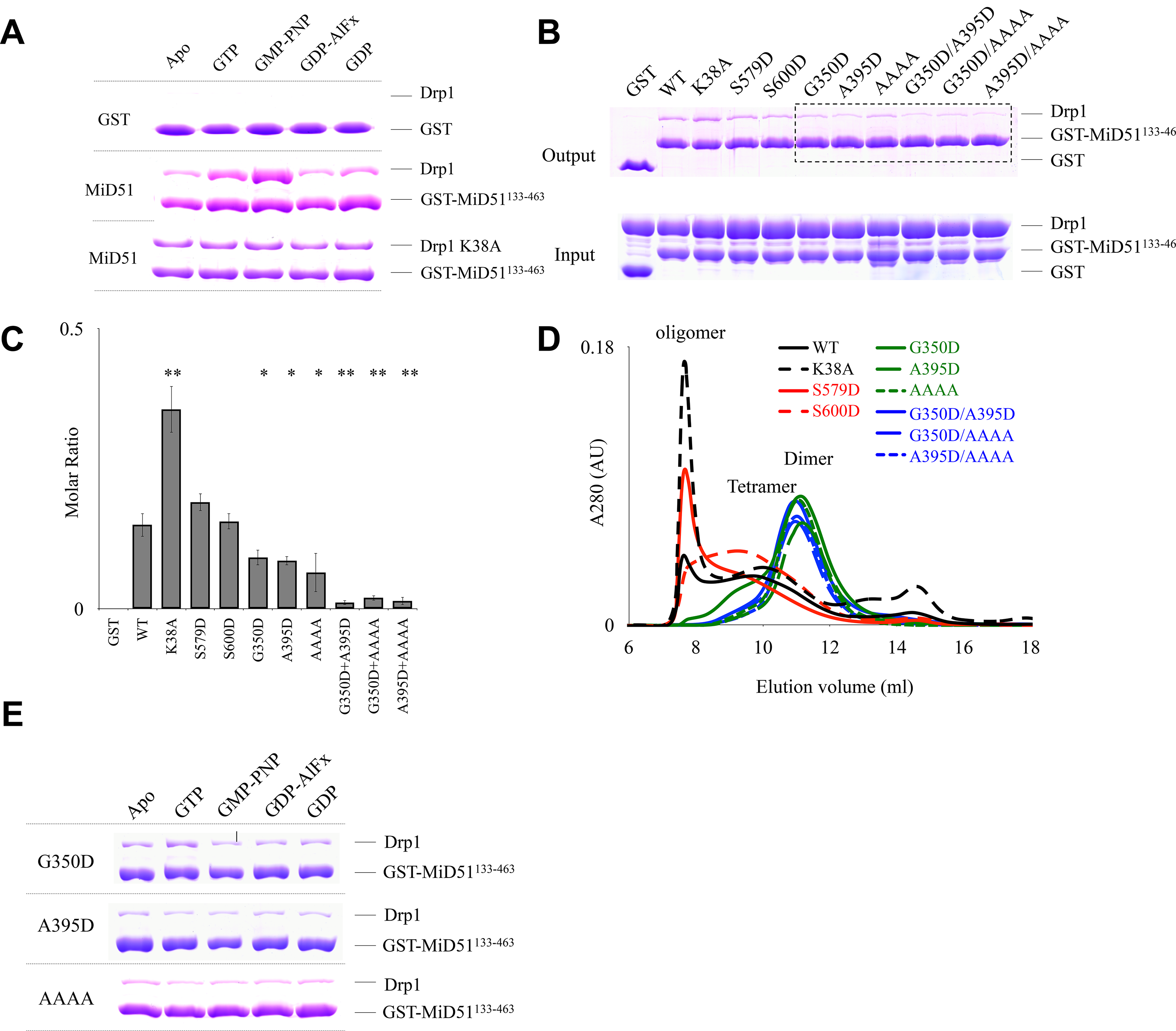
The cytoplasmic domain of MiD51 interacts with Drp1 is dependent on Drp1 oligomerization (A) Pull-down assays were performed to test the binding of purified Drp1 to GST-MiD51^133-463^ in the presence of different nucleotides. MiD51 and Drp1 or their mutants were mixed evenly before adding to the same amount of resin with the same volume to ensure equal amount of protein was used, and then 1 mM nucleotide at final concentration was added. (B) Pull-down assays were performed to demonstrate that the binding of Drp1 to MiD51 depends on Drp1 oligomerization. Purified GST, and GST-MiD51^133-463^ were loaded onto Glutathione Sepharose beads, and incubated with wild-type and mutated Drp1 to test their binding by SDS-PAGE. (C) Quantification of the results in (B). The binding affinity is expressed as molar ratio of Drp1 to MiD51 mutants. Data are shown as mean ± SEM of three independent experiments performed in triplicate, * P < 0.05; ** P < 0.005 compared to wild-type. (D) Gel filtration profiles of Drp1 and Drp1 mutants as indicated. Gel-filtration was performed with the size-exclusion column Superdex 200 PC 3.2/20 (GE Healthcare). The elution peak at ~11 ml represents the Drp1 dimer. (E) Pull-down assays were performed with purified Drp1 mutants in the presence of different nucleotides.

It is well established that Drp1 either in the presence of GTP or GMP-PNP forms oligomeric assemblies^19, 20^. To further clarify that whether the apparent greater binding between MiD51 and Drp1 in the presence of GTP or GMP-PNP relies on the oligomerization of Drp1, we made a series of Drp1 mutations and tested their effects on binding to MiD51. In previous studies ^21–25^, Drp1 residues G350, G363, R376, A395 and G401 play important roles in its polymerization. We designed a series of mutants where these residues were changed to Asp and monitored their ability to polymerize using gel filtration. We found that G350D and A395D behaved differently from wild-type Drp1 in gel filtration (Fig. 1D), suggesting defective oligomerization. Similar defects in oligomerization were also observed for the AAAA mutant form of Drp1 (^401^GPRP^404^→AAAA), ^23^. In addition, the compound mutants G350D/A395D, G350D/AAAA and A395D/AAAA showed a severe reduction in oligomerization. Our results indicate that the A395, G350 and GPRP (401-404) residues are involved in the polymerization of Drp1, consistent with previous studies. We next assessed the interactions between MiD51^133-463^ and Drp1 mutants. We found that compared to wild type, Drp1 oligomerization mutants G350D, A395D, AAAA, G350D/A395D, G350D/AAAA and A395D/AAAA have reduced affinity for MiD51^133-463^ (Fig. 1B), confirmed by quantification (Fig. 1C), and the mutant proteins generally have the same binding affinity for MiD51^133-463^ in the presence of different nucleotides (Fig. 1E). We also designed other Drp1 mutant proteins, targeting residues not responsible for oligomerization, such as K38A, and phosphorylation-mimic mutants S579D and S600D (related to S616 and S637 in Drp1 isoform 1). We found that these three mutant proteins behaved similarly to wild type protein based on both gel filtration and the interaction assay with MiD51 (Fig. 1B,C and D). These results indicate that the interaction between MiD51 and Drp1 significantly depends on Drp1 oligomerization.

### Structural analysis of the cytoplasmic domain of MiD51

To understand how MiD51 interacts with Drp1 during mitochondrial fission, we performed crystal structure studies and solved two types of MiD51 crystal structures under apo conditions (Table S1). Type I contains the cytosolic domain MiD51^129-463^, and the crystal space group is *P*4_1_2_1_2 with one molecule per asymmetric unit. Type II contains the fragment MiD51^133-463^, which was expressed as a C-terminal 6 × His fusion protein, and the crystal space group is *P*1 with two molecules per asymmetric unit. The overall structure consists of a central β-strand region flanked by two α-helical regions (Fig.2A) and looks similar to NTPase family crystal structures published by two groups ^17, 18^. The C domains of the Type I and Type II crystal structures are almost identical, but there is a tiny structural conformation change in the N domain (Fig. S1B), with a RMSD (root mean square deviation) variation of 1.14 Å for 329 aligned C_α_ atoms. By comparison, we found that all of the released crystal structures of MiD51 from PDB (Table S2), which include different nucleotide forms (Apo, ADP or GDP), lack distinct conformational changes when compared to the Type I and Type II crystal structures, with RMSD variations ranging from 0.97 to 1.88 Å (Fig. S1C and Table S3). We are not sure about the significance of such a small conformational change, which is probably due to different constructs, crystallization conditions and crystal packing. And the ADP/GDP binding sites are almost identical, which implies the structural rigidity of MiD51. It was reported that MiD51 can form dimers primarily based on the crystal packing of MiD51, but we did not observe such packing in our two crystal types (Fig. S1D). Further studies are needed to determine the oligomer state of MiD51, and we describe these studies in a later section.

### Two sites on MiD51 are involved in the interaction with Drp1

The crystal structure of MiD51 supplies limited information about Drp1 binding. Therefore, we performed a systematic analysis of MiD51 mutants (Table S4) to identify which region is involved in the interaction with Drp1. Initially we designed a series of mutant proteins, each containing a cluster of three or four mutated residues. We then used a pull-down assay to test the affinity of each MiD51 mutant for Drp1. These assays indicated that six MiD51 mutations disrupt the interaction with Drp1 (Fig. S2A and B). Next, we did a second round of point mutations of MiD51. We found eight mutant proteins with decreased affinity for Drp1 (Fig. 2C and D, Fig. S2C and D). There was almost no conformational change in the mutant proteins compared to wild type MiD51 based on circular dichroism (CD) spectroscopy and thermal shift assay (Fig. S2E and F).

**Figure 2.**
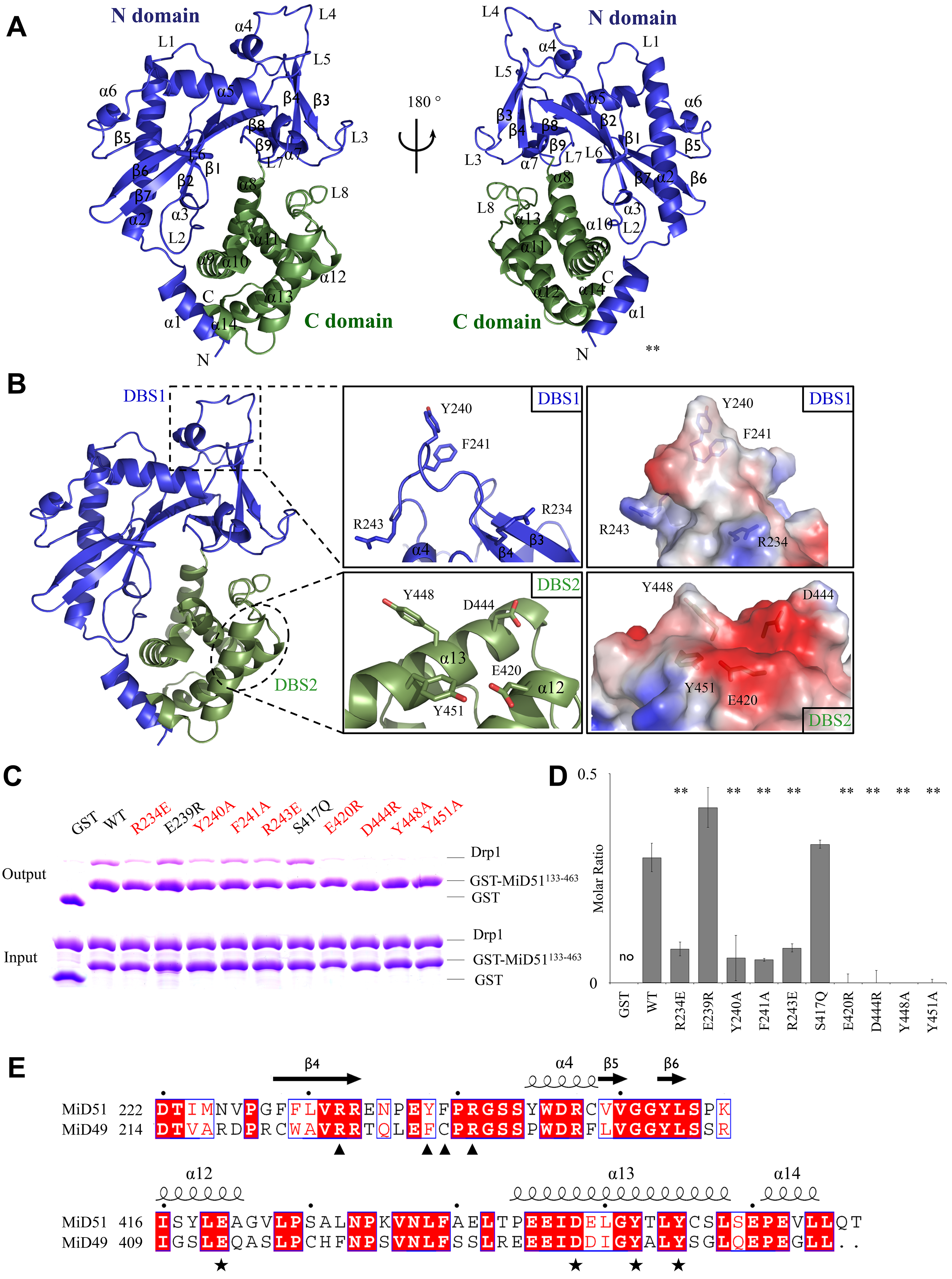
Two sites on MiD51 are involved in the interaction with Drp1 (A) Crystal structure of MiD51^133-463^. The N domain is colored blue, and the C domain is colored green. Secondary structure elements are labeled. (B) MiD51 sites that bind to Drp1. Left: Overview of the two MiD51 sites, which are outlined in dotted rectangles. Middle: Close-up views show the two binding sites. Key residues are labeled. Right: Electrostatic surface representation of two binding sites, with blue coloring indicating positive charges and red coloring indicating negative charges. (C) WT and mutant GST-MiD51^133-463^ in vitro pull-down assays were performed with purified Drp1. (D) Quantitation of the results in (C). The binding affinity is expressed as molar ratio of Drp1 to MiD51 mutants. Data are shown as mean ± SEM of three independent experiments performed in triplicate, with ** P < 0.005 compared to wild-type. (E) Sequence alignment of MiD51 and MiD49 sequences. Strictly conserved residues are highlighted in red. Secondary structural elements are depicted on the top of the alignments. Residues involved in Drp1 interaction are marked with★ for DBS1 and ▴ for DBS2.

We analyzed the distribution of these sites and found that the eight mutations are located in two areas. The first area contains four residues, R234, Y240, F241 and R243, which are located on an exposed loop between β4-α4 (Fig. 2B). When these residues are substituted with alanine or glutamate (R234E, Y240A, F241A and R243E), the resulting mutant proteins have modest or serious decreases in Drp1 binding affinity, as confirmed by quantitation (Fig. 2C and D). This suggests that the exposed loop is a main determinant for Drp1 binding. We name this area DBS1 (Drp1 Binding Site One) (Fig. 2B), which is consistent with previous studies ^17, 18^. The second area contains the amino acids E420, D444, Y448 and Y451(Fig. 2B). Mutation of these residues by substituting with alanine, or by substituting aspartate and glutamate with arginine (E420R, D444R, Y448A and Y451A), results in a more dramatic effect on the ability of MiD51 to bind Drp1, and in some cases even abolishes binding (Fig. 2C and D). We define this area as DBS2 (Drp1 Binding Site Two), which is located on α12 and α13 in the C domain and forms a surface for Drp1 binding (Fig. 2B). Therefore, MiD51 requires DBS2, a surface in the C domain, to cooperate with Drp1 binding. An amino acid sequence alignment of MiD51 and MiD49 proteins from different species reveals that these eight DBS1 and DBS2 residues are highly conserved (Fig. 2E and Fig. S2G). Based on the crystal structure of MiD49 ^18^, these eight residues also form an exposed loop in the N domain and a surface in the C domain for Drp1 binding.

### MiD51 forms a dimer via an disulfide bond and is important for interaction with Drp1

Mitochondrial fission receptors, such as Fis1 and Mff, form dimers to perform their functions in mitochondrial fission ^12^. A previous study reported that MiD51 could form a dimer under non-reducing conditions ^26^, suggesting that MiD51 may form a dimer via an intermolecular disulfide bond between cysteines in the region of residues 49 to 195 ^26^. But based on the crystal structure, another study found that MiD51 forms a dimer via electrostatic interactions in the N-terminal helix, and the dimerization is very important for its function in mitochondrial fission^18^. Surprisingly, we did not observe a similar surface mediating the dimerization of MiD51 in our crystal packing. Therefore we experimentally determined whether MiD51 forms dimers. Using a time course assay where the level of dimer formation was quantified every twenty-four hours, we determined that MiD51^133-463^ does form dimers and that the level of dimerization continues to increase over time (Fig. 3A and B). These results correlate well with the results of Zhao et al ^26^. However, a limited amount of MiD51^133-463^ protein exists as dimers based on native-PAGE and gel filtration experiments (Fig. 3C and D), indicating that the majority of MiD51^133-463^ exists as a monomer.

**Figure 3.**
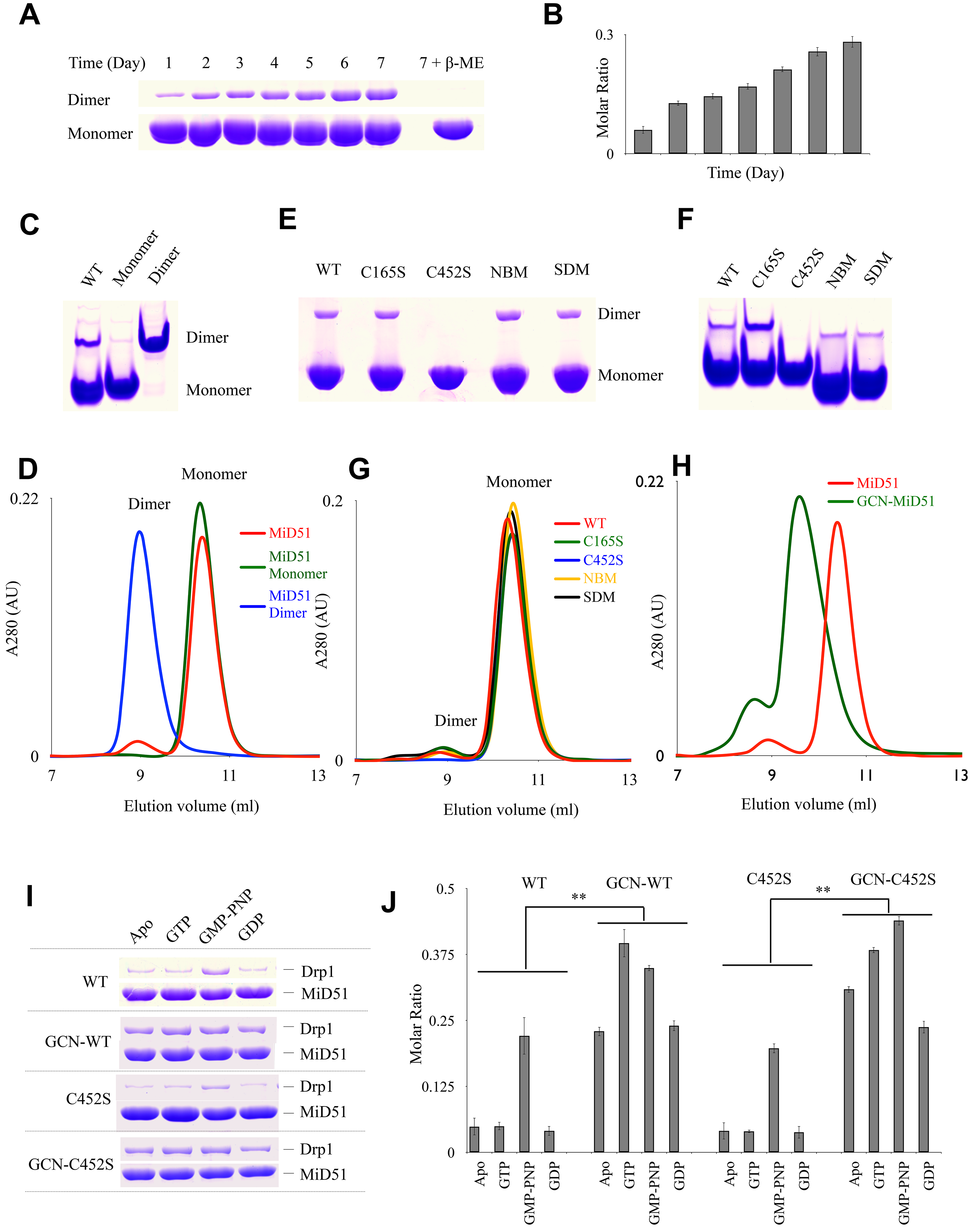
MiD51^133-463^ forms a dimer via an intermolecular disulfide bond between C452 residues and is important for its interaction with Drp1. (A) A time course experiment, where the level of dimer formation was quantified every twenty four hours, and non-reducing SDS-PAGE indicates that MiD51^133-463^ can form dimers in the air. (B) Quantification of the results in (A). The level of dimerization is expressed as the ratio of dimer to monomer. All error bars represent SD from three independent experiments. (C) Native PAGE analysis of monomeric and dimeric MiD51^133-463^. (D) Gel filtration analysis of monomeric and dimeric MiD51^133-463^. Gel-filtration was performed with the size-exclusion column Superdex 75 PC 3.2/20 (GE Healthcare). The blue profile represents dimer and the green profile represents monomer. (E), Non-reducing SDS-PAGE of wild-type and mutant MiD51^133-463^ shows that the C452S mutant is not able to form dimers. (F) Native PAGE of wild-type and mutant MiD51^133-463^ also indicates that the C452S MiD51^133-463^ mutant does not form dimers. (G) Gel filtration analysis of MiD51^133-463^ and mutants shows that the dimer peak is missing in the C452S mutant. (H) Gel filtration profiles illustrate the increased dimerization of the dimer-mimic GCN-MiD51, compared to wild type MiD51. (I) Strep pull down of MiD51, GCN-MiD51, C452S, and GCN-C452S proteins with Drp1. Equal amount of purified wild-type and mutant MiD51^133-463^ proteins were loaded onto 15 μl Strep-Tactin resin, and incubated with Drp1 to test its binding in the presence of different nucleotides by SDS-PAGE. The GCN peptide fusion was used to obtain artificial MiD51 dimers. (J) Quantification of the results in (I). The binding affinity is expressed as molar ratio of Drp1 to MiD51 mutants. Data are shown as mean ± SEM of three independent experiments performed in triplicate, ** P < 0.005.

To determine which residue mediates MiD51 dimerization, we analyzed the MiD51 sequence and found that there are seven cysteines, but only two, C165 and C452, are exposed on the protein surface. When C452 was substituted by serine (C452S) the MiD51 protein lost its ability to form dimers, as shown by lack of dimer formation in non-reducing PAGE, Native PAGE and gel filtration experiments (Fig. 3E, F and G). In contrast, another cysteine mutant C165S showed dimerization similar to wild type (Fig. 3E, F and G). This suggests that C452, not C165, is the residue that forms the disulfide bond. We also found that the supposed dimer surface mutant (SDM, R169A/R182A/D183A/Q212A/N213A) and nucleotide binding site mutant (NBM, H201D/R342E/K368E) (Fig. S1A) were still capable of dimerization similar to wild type (Fig. 3E, F and G). Therefore, the dimerization of MiD51 is not mediated by the previously supposed dimer surface^18^ or the nucleotide binding site. Thus, the results certify that MiD51 forms dimers via an intermolecular disulfide bond between C452 residues.

We next asked if the dimerization of MiD51 plays a critical role in Drp1 binding. Considering that the percentage of MiD51 that exists as a dimer is low in *E. coli* that overexpress MiD51, we added a GCN peptide (ARMKQLEDKIEELLSKIYHLENEIARLKKLIGER), which forms a dimer ^27^, to the amino terminal end of MiD51 to make an artificial MiD51 dimer (Fig. 3H). We then checked the interaction between MiD51 and Drp1 under different nucleotide acid binding states. Strep pull-down assays indicated that wild-type Drp1 interacts with MiD51^133-463^ in a weak manner, although the GMP-PNP bound state had a higher affinity than other states, including the Apo, GTP and GDP states. When expressed as a GCN peptide fusion, MiD51^133-463^ had increased binding affinity for Drp1 regardless of the nucleotide binding state (Fig. 3I and J). Even C452S mutant protein showed the same binding affinity for Drp1 as wild type, when expressed as a GCN peptide fusion protein (Fig. 3I and J). These results indicate that dimerization of MiD51 enhances its binding affinity for Drp1.

## DISCUSSION

The role of MiD51 in mitochondrial fission has been well established ^15,17,26,28–30^. MiD51 mediates mitochondrial fission by recruiting Drp1 to the outer mitochondrial membrane and regulating its assembly and mitochondrial fission activity in a GTP-dependent manner. We have elucidated in molecular detail how the interaction between MiD51 and Drp1 is regulated by multi-faceted mechanism.

The changes in Drp1 conformation and oligomerization upon GTP binding, hydrolysis and release, is associated with the procession of mitochondrial fission ^31^. We suggest that the interaction between MiD51 and Drp1 undergoes changes during this process. Initial insight came from the binding of MiD51 for Drp1 under different nucleotide binding states. We determined that MiD51 binds effectively to the GTP and GMP-PNP bound states of Drp1, which depends on the polymerization of Drp1. Recent research suggests that oligomerization of Drp1 is required for its interaction with Mff, whereas MiD51 does not have a strong requirement for Drp1 oligomerization because AAAA mutant form of Drp1, which are only capable of forming dimers, still show binding activity for MiD51 ^32^. Although the AAAA mutant has the capacity to bind MiD51, its binding affinity is reduced compared to wild type as we showed (Fig. 1C and D).

We then gained significant insight into the interaction between MiD51 and Drp1 by examining the contact interface on MiD51. By performing a systematic screen of proteins with mutations in surface residues, we determined that two regions, DBS1 and DBS2, in MiD51 make direct contact with Drp1. DBS1, containing R234, Y240, F241 and R243, is located on an exposed loop of the N domain. The location of this binding site is consistent with previous studies ^17^, one of which proposed that the topology of the loop is a critical factor for Drp1 binding. This suggests that electrostatic interactions and hydrophobic interactions may play important roles in the binding of Drp1 to MiD51. DBS2 is a novel region located on the surface of the C domain. Single mutations, such as E420R, D444R, Y448A and Y451A, completely abolish MiD51-mediated binding of Drp1, suggesting that DBS2 is much more important than DBS1 for Drp1 recruitment. We know that the interaction between MiD51 and Drp1 changes during the process of mitochondrial fission, so the interaction may need more than one binding site between MiD51 and Drp1. In addition to the exposed loop in N domain, we have determined that another region in MiD51 makes direct contact with Drp1, and the residues are highly conserved between MiD51 and MiD49. It seems likely that the two binding regions on MiD51 are responsible for the complicated interaction with Drp1 during the process of mitochondrial fission. But the precise role of MiD51 in Drp1 polymerization and mitochondrial fission still remains elusive.

We also determined that MiD51 forms dimers via an intermolecular disulfide bond between C452 residues located in the C terminal region, although the majority of MiD51 protein exists as monomer. The monomer-dimer state of MiD51 is closely related to its interaction with Drp1 in mitochondrial fission because the interaction between MiD51 and Drp1 is enhanced by dimerization of MiD51. This could be reflected coincidently by pull-down assays, in which the binding affinity of monomeric Strep-tagged MiD51 for Drp1 is weaker than that of dimeric GST-fused MiD51 with Drp1, especially at GTP bound state (Fig. 1A and 3I). So the mitochondrial fission activity of Drp1 could be regulated by the metabolism state of cells through increased interaction with dimerized MiD51. We note that the Drp1 receptor Mff exits as a tetramer formed via its coiled coil region, and only multimeric Mff can bind Drp1 effectively, facilitate assembly of Drp1 polymer, stimulate GTPase activity and trigger mitochondrial fission ^18, 33^. So the MiD51 and Mff receptors function in a similar way by forming a dimer or tetramer to recruit Drp1 and regulate mitochondrial fission. However, the C452 residue required for dimerization of MiD51 is not conserved in MiD49, so MiD51 and MiD49 might mediate mitochondrial fission via different regulatory mechanisms. Further work will be necessary to understand whether and how MiD49 forms dimers to regulate fission.

Collectively, we propose a model for MiD51-mediated recruitment of Drp1 to regulate mitochondrial fission (Fig. 4). **1.** At basic conditions, most Drp1 protein is inactive in the cytoplasm, and MiD51 doesn’t form dimers; therefore, only a small amount of Drp1 binds to MiD51; 2. For mitochondrial fission, Drp1 binds to GTP and undergoes oligomerization and MiD51 forms dimers via disulfide bond formation between C452 residues, leading to the enhancement of the interaction between MiD51 and Drp1 by DBS1 and DBS2; 3. Dimeric MiD51 recruits more oligomeric Drp1 to the mitochondrial outer membrane, resulting in the formation of the mitochondrial fission complex around the fission site; 4. GTP hydrolysis further enhances the interaction between MiD51 and Drp1, and triggers mitochondrial fission by the fission complex and other factors such as Dyn2 and endoplasmic reticulum; 5. After mitochondrial fission is complete along with the production of GDP, oligomeric Drp1 depolymerizes, the interaction between MiD51 and Drp1 is weakened to that observed at basic levels, and finally Drp1 is released from the membrane and localizes to cytoplasm where it is free to function in another cycle of mitochondrial fission.

**Figure 4.**
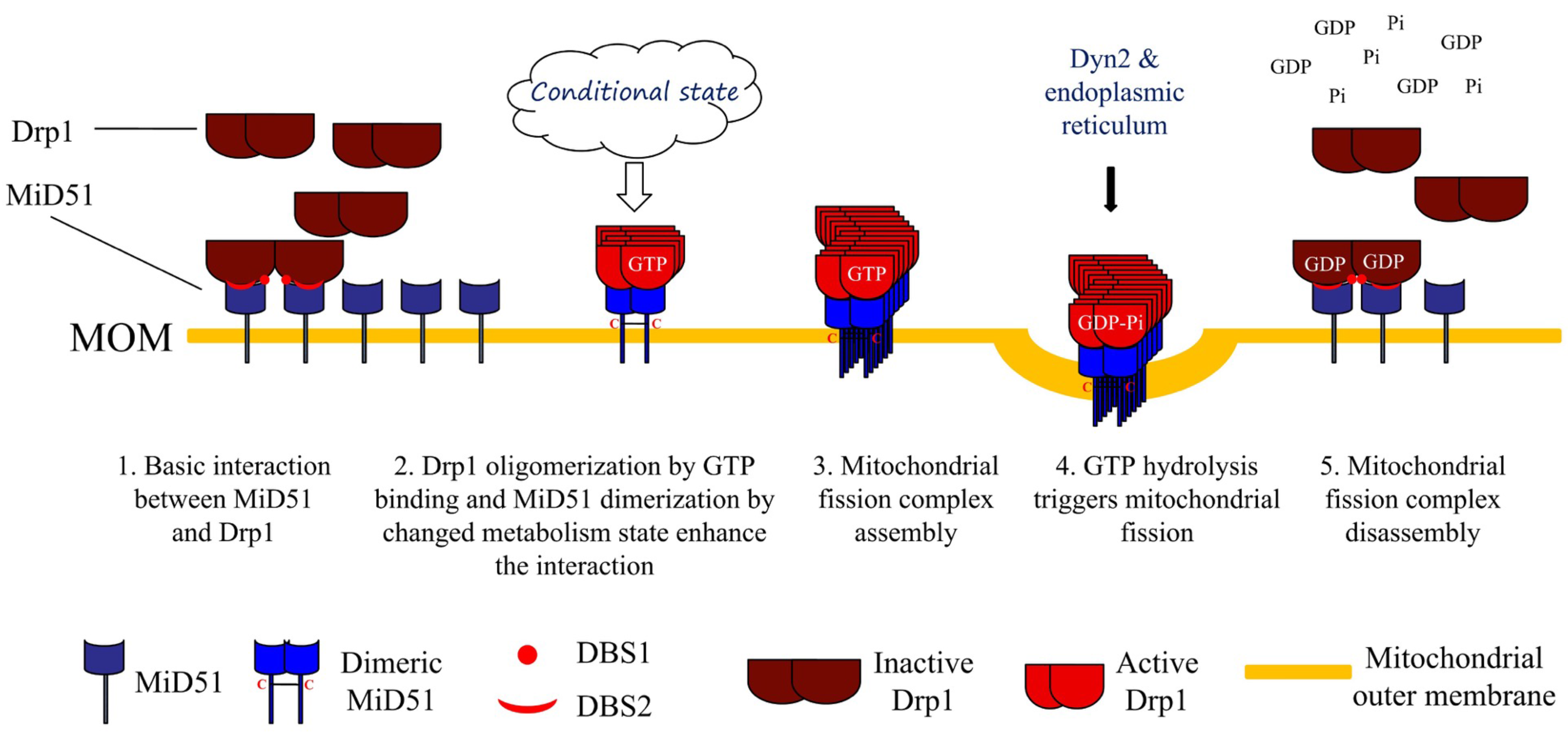
A proposed model for MiD51-mediated recruitment of Drp1 and mitochondrial fission.

## MATERIALS AND METHODS

### Molecular cloning and plasmid constructions

The open reading frames (ORF) encoding MiD51^ΔTM^ MiD51^129-463^, MiD51^133-463^ and mutants were amplified by PCR from the full-length human MiD51 ORF (GenBank accession No. NM_019008) and cloned into the pGEX-6P-1 vector (GE Healthcare), or pET22b (Novagen) derivative vector with a N-terminal 6*His tag or Strep tag. Drp1 (GenBank accession No. NM_005690) and its mutants were cloned into the pET22b (Novagen) derivative vector with a C-terminal 6*His tag or Strep tag. All site-directed mutagenesis of MiD51 and Drp1 were performed by the overlapping PCR method.

### Protein expression and purification

All constructs of MiD51 were expressed in *Escherichia coli* BL21 (DE3) (Invitrogen). Recombinant proteins were induced by addition of 0.3 mM IPTG at a culture density of OD600~0.6, followed by 16 h incubation at 16°C. To purify His tag proteins, the bacterial cells were lysed by high-pressure homogenization in lysis buffer containing 50 mM Tris-HCl, 300 mM NaCl, 40 mM imidazole, pH 8.0.

Cell debris was removed by centrifugation at 16,000 g for 40 min, the supernatant was applied to Ni-NTA Sepharose (GE Healthcare) and washed with lysis buffer, and the protein was eluted with lysis buffer plus 300 mM imidazole and concentrated using Amicon Ultra-4 centrifugal filter units (10 kDa cutoff, Millipore). A HiTrap Desalting column (5 ml, GE Healthcare) was used to change the buffer of proteins to 20 mM Tris-HCl pH 8.0, 50 mM NaCl. The protein was further purified by anion-exchange chromatography on a Resource Q column (GE Healthcare) with a NaCl linear gradient of 50-600 mM in 20 mM Tris-HCl pH 8.0. The eluted fractions containing MiD51 were pooled and concentrated, and finally purified by size-exclusion chromatography using a Superdex 75 column (GE Healthcare) pre-equilibrated with 20 mM HEPES pH 7.5, 150 mM NaCl. To purify strep tag proteins, the bacterial cells were lysed in buffer containing 10 mM Na2HPO4, 1.8 mM KH2PO4, 140 mM NaCl, 2.7 mM KCl, 1 mM DTT pH 7.4. The supernatant was applied to Strep-Tactin Sepharose (IBA GmbH) and the protein was eluted with lysis buffer plus 2.5 mM desthiobiotin. The other steps were the same as His tag proteins.

To purify GST fusion proteins, bacterial cells were lysed in lysis buffer (10 mM Na2HPO4, 1.8 mM KH2PO4, 140 mM NaCl, 2.7 mM KCl, 1 mM DTT pH 7.4), and the supernatant was applied to Glutathione Sepharose (GE Healthcare) loaded into a 20-ml gravity flow column. For crystallization, the resins were first washed with the lysis buffer, then the GST fusion proteins were digested using PreScission protease (GE Healthcare) at 16°C overnight on column. The digested MiD51 protein was eluted using the lysis buffer. The next steps were the same as His tag proteins. For GST pull-down assaay, GST-fused proteins were directly eluted with 20 mM reduced glutathione, 50 mM Tris-HCl pH 8.0, 150 mM NaCl. After concentration, GST-fused protein was directly changed the buffer to 20 mM HEPES pH 7.5, 150 mM NaCl. The selenomethionine (SeMet)-labeled MiD51^129-463^ was prepared as described previously ^34, 35^. In brief, the expression vector containin GST-fused MiD51^129-463^ was transformed into the methionine auxotroph *E. coli* B834 strain (Novagen). The cells were grown in M9 medium supplemented with YNB medium, 50g/L glucose, 2mM MgSO4, 0.1 mM CaCl2 and 50 mg/L of L-selenomethionine. The purification process of the SeMet-labeled MiD51^129-463^ was the same as that used for the native protein.

Wild type Drp1 and its mutants were expressed in *E. coli* Rosseta (DE3) cells (Invitrogen), and the expression and purification process was in the same way as MiD51^133-463^ with 6*His tag or strep tag respectively.

### GST and strep pull-down assay

For GST pull-down assay, equal amounts of GST, GST-fused MiD51^133-463^, and GST-fused mutant proteins were loaded onto 15 μl of Glutathione Sepharose 4B slurry beads in assay buffer (20 mM HEPES, pH 7.5, 150 mM NaCl, 1% Triton X-100, 1mM MgCl_2_). After incubation with equal molar of Drp1 for 3 h at 4°C, the pellets were washed three times with 500 μl of assay buffer, subsequently incubated with SDS-PAGE sample buffer at 95°C, separated on 12% SDS-PAGE gels and detected using Coomassie blue staining. In the case of pull-down assay with different nucleotides (ATP, AMP-PNP, ADP-AlFx or ADP), MiD51 and Drp1 were mixed evenly first before dispensing the same amount of volume to the same amount of resin, and then 1 mM nucleotide at final concentration was added in the assay buffer and wash buffer. For strep pull-down assay, equal amounts of strep-fused wild type and mutant MiD51 proteins were loaded onto 15 ul of Strep-Tactin Sepharose (IBA GmbH). The following steps are same as the GST pull-down assay.

### Crystallography

Crystals of MiD51^133-463^ and Se-MiD51^129-463^ were obtained using the hanging drop vapor diffusion method at 16°C. To set up a hanging drop, 1 μl of concentrated protein solution was mixed with 1 μl of crystallization solution. The final optimized crystallization condition was 0.6 M NaH_2_PO_4_/K_2_HPO_4_, pH 7.0 for MiD51^133-463^ at 30mg/ml and 0.2 M L-Proline, 0.1 M HEPES pH 7.0, 6% PEG3350 for MiD51^129-463^ at 18mg/ml. Before X-ray diffraction, crystals were soaked in crystallization solution containing 20% glycerol for cryo-protection. The diffraction data for native and SeMet derivative crystals were collected at 100 K at beamline BL17U at Shanghai Synchrotron Radiation Facility (SSRF). The diffraction data were processed and scaled using HKL2000 ^36^. The structure of MiD51^129-463^ was solved with the single-wavelength anomalous diffraction (SAD) method. Selenium atoms were successfully located with the SHELXD ^37^ in HKL2MAP ^38^. Phases were calculated and refined with SOLVE and RESOLVE ^39, 40^. An initial model was built using COOT ^41^ and further refined using REFMAC5 ^42^. The structure of MiD51^133-463^ was solved by molecular replacement with Phaser ^43^ and further refined using REFMAC5. The stereo-chemical quality of the final model was validated by PROCHECK ^44^ and MolProbity ^45^. The statistics for data processing and structure refinements are listed in Table S1. All structural figures were prepared with PyMOL (http://www.pymol.org/).

## ADDITIONAL INFORMATION

**Competing Interests:** The authors declare that they have no competing interests.

**Accession codes:** The coordinates for the crystal structures of MiD51^129-463^ and MiD51^133-463^ have been deposited in the Protein Data Bank (PDB), with the accession codes 5X9B and 5X9C respectively.

## AUTHOR CONTRIBUTIONS

F. S. and Q.C. initiated the project. J.M. and F.S. designed all the experiments. J.M., Y.Z., K.Z., M.C., and X.P. performed the experiments. J.M., X.P. and F.S. analyzed the data and wrote the manuscript.

## ACKNOWLEDGMENTS

We are grateful to Ruigang Su and Ping Shan (Fei Sun’s group) for assistance in lab management. This project is financially supported by the National 973 Program from the Chinese Ministry of Science and Technology (Grants 2014CB910700 and 2011CB910301), and the Strategic Priority Research Program (XDB08030202) from the Chinese Academy of Sciences (CAS).

## SUPPLEMENTAL MATERIALS

### EXPERIMENTAL PROCEDURES

#### Thermal shift assay and circular dichroism (CD) spectroscopy

For thermal shift assay, the wild type and mutant 6 × His-MiD51^133-463^ protein samples were diluted to 1 mg/ml in the buffer containing 20 mM HEPES pH 7.5, 150 mM NaCl. The fluorescent dye SYPRO Orange (Invitrogen) was added into the sample solution by ~1,000 fold of dilution. 20 μl of mixture in a PCR tube was heated up from 25°C to 75°C with the step of 1°C per min. The fluorescence of the mixture was measured by using a RT-PCR device (Corbett 6600). The melting temperature (Tm) was estimated as the temperature corresponding to the minimum of the first derivative of the protein denaturation curve. All the measurements were repeated three times.

For CD spectroscopy assay, the wild type and mutant 6XHis-MiD51^133-463^ protein samples were diluted to 0.2 mg/ml in buffer containing 10 mM Na_2_HPO4, 1.8 mM KH_2_PO4, 140 mM NaCl, 2.7 mM KCl, 1 mM DTT, pH 7.4. The spectra were recorded over the wavelength from 200 nm to 260 nm with a bandwidth of 1 nm and 0.5 s per step by using CD spectrometer (Chirascan-plus, Applied photphysics). All the measurements were repeated three times and the spectrum data were corrected by subtracting the buffer control.

**Fig. S1.**
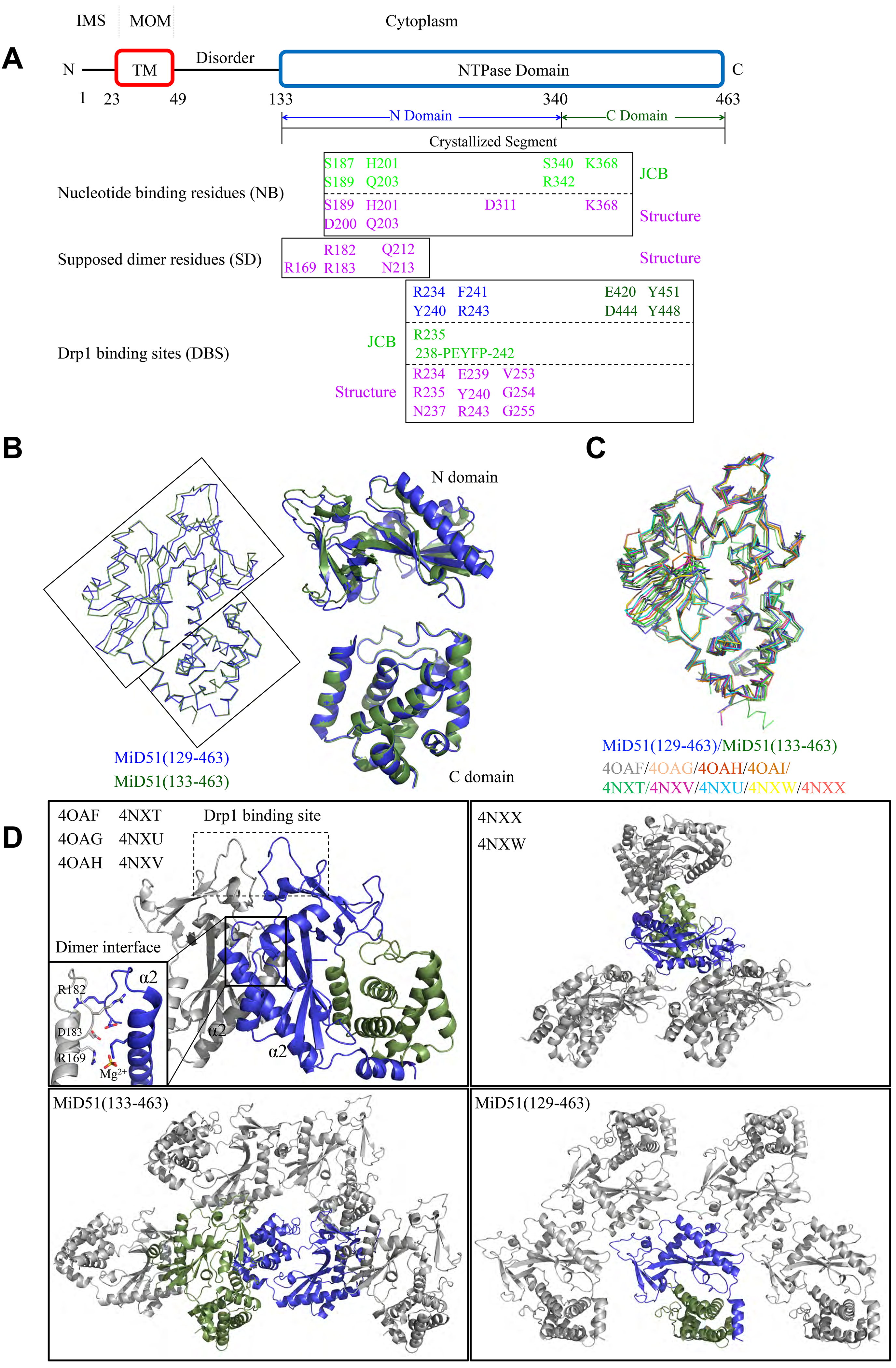
Comparisons between Mid51 protein crystal structures. () Topology of MiD51 and key residues involved in nucleotide binding, dimerization and Drp1 binding. Domain boundaries are marked with residue numbers. The NTPase domain can be divided into two sub-domains, N domain (133-339) and C domain (340-463). TM, transmembrane domain; IMS, inter-membrane space; MOM, mitochondrial outer membrane. (B) Comparison of the cytoplasmic domain in crystal structures of MiD51, Mid51^129-463^ and Mid51^133-463^. (C) Comparison of the crystal structure of MiD51^133-463^ with the cytoplasmic domain crystal structure of MiD51 from PDB (codes 4OAF, 4OAG, 4OAH, 4NXT, 4NXV, 4NXU, 4NXW and 129 463 4NXX), and comparison of the crystal structure of MiD51^129-463^ with the crystal structure of MiD51 from PDB (code 4OAI). (D) Crystal packing of the MiD51 structures shown in (C).

**Fig. S2.**
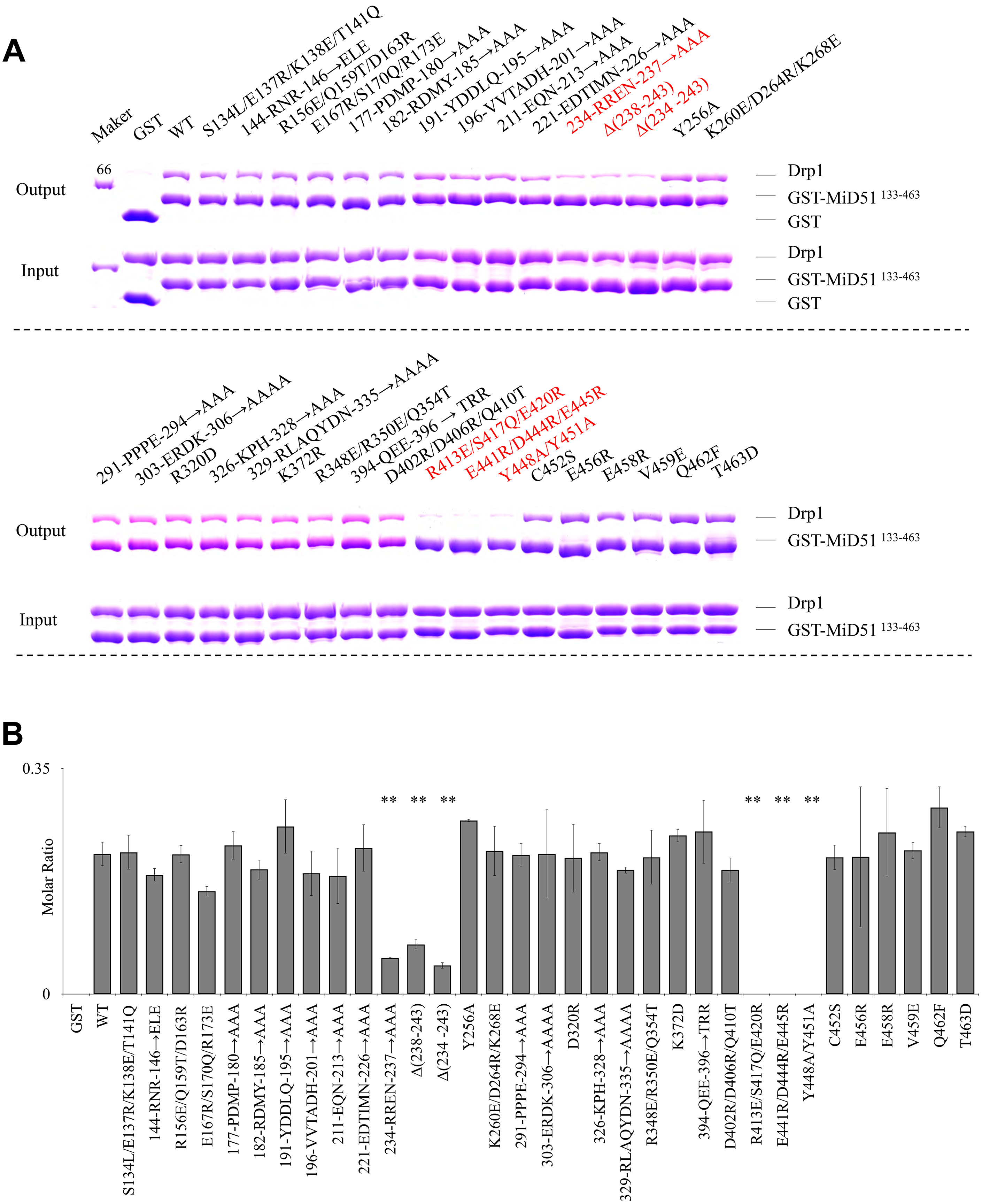
Systematic mutation screening to investigate the regions of Mid51 that are involved in the interaction with Drp1. (A) Mutant forms of MiD51 containing clusters of three or four mutated residues were initially tested for ability to bind Drp1 with in vitro GST pull-down assays. Six MiD51 mutants that disrupt the interaction with Drp1 are colored in red. (B) Quantification of the results in (A). The binding affinity is expressed as molar ratio of Drp1 to MiD51 mutants. Data are shown as mean ± SEM of three independent experiments performed in triplicate, with ** P < 0.005 compared to wild-type. (C) In vitro GST pull-down assays were used to screen the single point mutants based on the results of (A) and (B). Mutations that disrupt the interaction with Drp1 are colored in red. (D) Quantification of the results in (C). The binding affinity is expressed as molar ratio of Drp1 to MiD51 mutants. Data are shown as mean ± SEM of three independent experiments performed in triplicate, with ** P < 0.005 compared to wild-type. (E) Circular dichroism spectroscopy confirmed that MiD51 mutants that have disrupted interactions with Drp1 still have the same conformation as wild type. (F) Thermal shift stability assays confirmed that there is not much conformational change in mutants compared to the wild type. (G) Sequence alignment of full-length MiD51 and MiD49 proteins. MiD51 and MiD49 proteins are distinguished by grey shading. Strictly conserved residues are highlighted in red, and moderately conserved residues are outlined in blue. Residues involved in Drp1 interaction are marked with ★ for DBS1 and▴ for DBS2. The secondary structures are shown above the sequences.

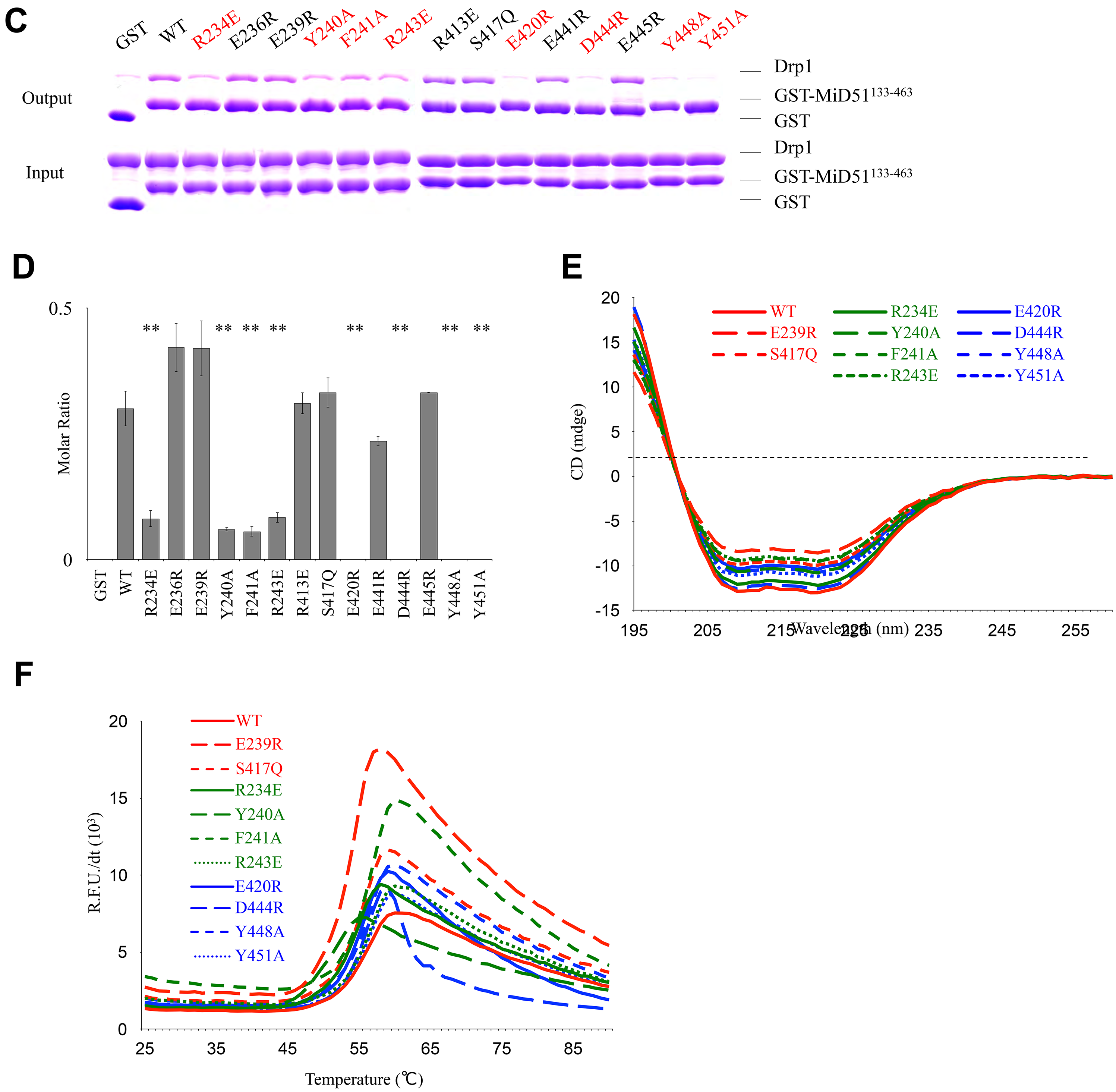

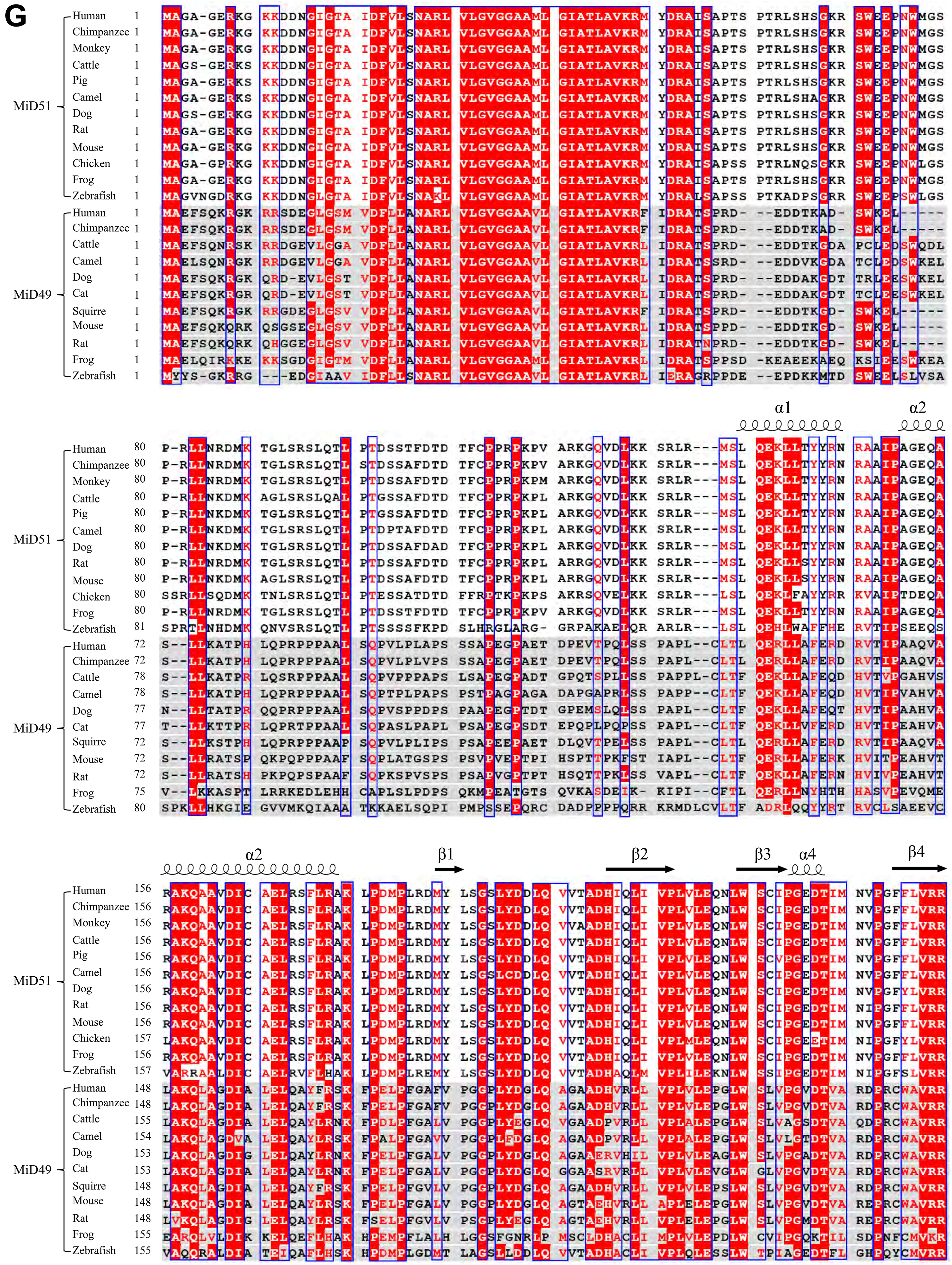

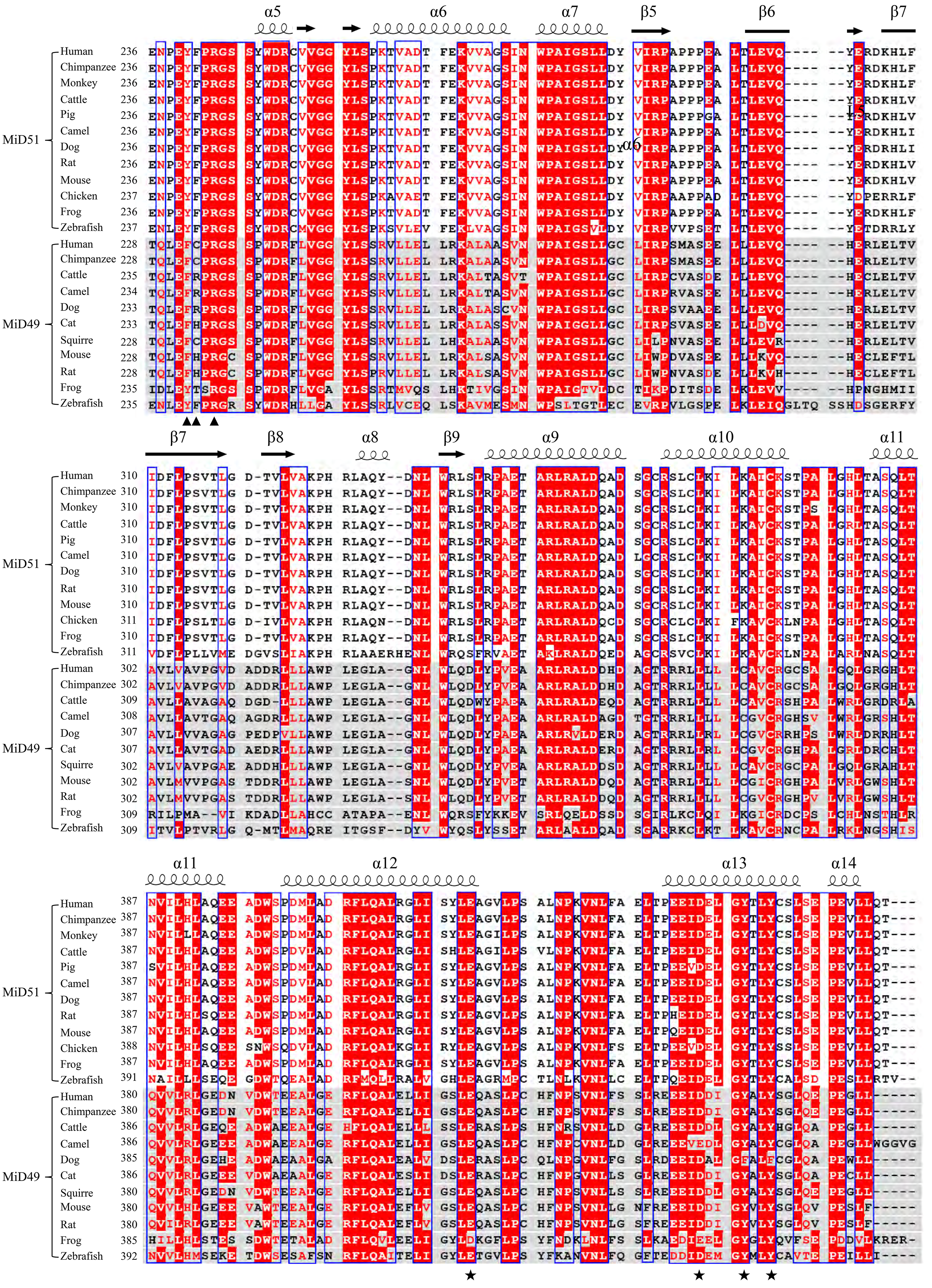

**Table S1.** Data collection and refinement statistics.

| Crystal | Se-MiD51<sup>129-463</sup> | MiD51<sup>133-463</sup> |
| --- | --- | --- |
| **Data collection** |  |  |
| Wavelength(Å) | 0.9793 | 0.9793 |
| Space group | P4<sub>1</sub>2<sub>1</sub>2 | P1 |
| Cell dimensions |  |  |
| *a*, *b*, *c* (Å) | 88.8, 88.8, 124.7 | 61.3, 64.7, 65.9 |
| $\alpha$, $\beta$, $\gamma$ (°) | 90.0, 90.0, 90.0 | 89.8, 108.1, 117.2 |
| Resolution (Å) | 50.00-2.70(2.80-2.70) | 50.00-1.85(1.92-1.85) |
| *R*<sub>sym</sub> (%) | 0.087(0.832) | 0.068(0.595) |
| *I*/σ*I* | 15.9(2.1) | 23.9(5.4) |
| Completeness (%) | 99.6 | 95.3 |
| Redundancy | 4.4 | 5.5 |
| **Refinement** |  |  |
| Resolution (Å) | 44.42-2.70(2.80-2.70) | 48.54-1.85(1.92-1.85) |
| No. reflections | 14284 | 68988 |
| *R*<sub>work</sub> / *R*<sub>free</sub> (%) | 0.227/0.251 | 0.203/0.232 |
| **No. atoms** |  |  |
| Protein | 2592 | 5168 |
| Water | 10 | 202 |
| **B factors (Å<sup>2</sup>)** |  |  |
| Protein | 84.92 | 22.83 |
| Water | 71.01 | 27.94 |
| **r.m.s. deviations** |  |  |
| Bond lengths (Å) | 0.015 | 0.022 |
| Bond angles (°) | 2.08 | 2.16 |

**Table S2.** Sum of partial crystallographic statistics for MiD51^129-463^, MiD51 ^133-463^, and released PDB crystal structures.

| PDB | Fragment | Crystal types | Space group | Unit-cell parameters | Molecules per asymmetric unit | Resolution | Reference |
| --- | --- | --- | --- | --- | --- | --- | --- |
| 5X5B | 129-463 | Native | $P4_12_12$ | 88.8, 88.8, 124.7, 90.0, 90.0, 90.0 | 1 | 2.70 | This study |
| 5X5C | 133-463 | Native | $P1$ | 61.3, 64.7, 65.9 89.8, 108.1, 117.2 | 2 | 1.85 | This study |
| 4OAF | 134-463 | Native | $P2_1$ | 91.1, 78.6, 102.3 90.0, 96.6, 90.0 | 4 | 2.20 | (Loson et al., 2014) |
| 4OAG | | ADP bound | $P2_1$ | 62.1, 80.8, 65.2 90.0, 105.7, 90.0 | 2 | 2.00 | (Loson et al., 2014) |
| 4OAH | | H201A | $P2_1$ | 82.4, 79.2, 103.5 90.0, 98.0, 90.0 | 4 | 2.00 | (Loson et al., 2014) |
| 4OAI | | CDM | $P2_12_12_1$ | 63.7, 67.1, 79.4 90.0, 90.0, 90.0 | 1 | 2.00 | (Loson et al., 2014) |
| 4NXT | 119-463 | Native | $P1$ | 72.7, 78.7, 79.4 66.3, 84.9, 64.1 | 4 | 2.12 | (Richter et al., 2014) |
| 4NXV | | GDP bound | $P1$ | 72.3, 79.1, 80.1 65.8, 84.4, 64.1 | 4 | 2.30 | (Richter et al., 2014) |
| 4NXU | | ADP bound | $P1$ | 72.6, 79.3, 79.4 65.4, 84.2, 63.4 | 4 | 2.30 | (Richter et al., 2014) |
| 4NXX | | GDP bound | $P4_32_12$ | 57.7, 57.7, 255.4 90.0, 90.0, 90.0 | 1 | 2.55 | (Richter et al., 2014) |
| 4NXW | | ADP bound | $P4_32_12$ | 57.7, 57.7, 253.8 90.0, 90.0, 90.0 | 1 | 2.55 | (Richter et al., 2014) |

**Table S3.** RMSD variations for superimposition of the Ca backbone of MiD51^129-463^, MiD51^133-463^, and released PDB crystal structures.

| RMSD |  | 1 | 2 | 3 | 4 | 5 | 6 | 7 | 8 | 9 | 10 |
| --- | --- | --- | --- | --- | --- | --- | --- | --- | --- | --- | --- |
|  |  | 4NXT | 4NXU | 4NXV | 4NXW | 4NXX | 4OAF | 4OAG | 4OAH | 4OAI | M129 |
| 2 | 4NXU | 1.13 |  |  |  |  |  |  |  |  |  |
| 3 | 4NXV | 0.91 | 0.38 |  |  |  |  |  |  |  |  |
| 4 | 4NXW | 1.40 | 1.13 | 1.14 |  |  |  |  |  |  |  |
| 5 | 4NXX | 1.40 | 1.07 | 1.18 | 0.18 |  |  |  |  |  |  |
| 6 | 4OAF | 0.54 | 1.19 | 0.98 | 1.45 | 1.45 |  |  |  |  |  |
| 7 | 4OAG | 0.94 | 1.09 | 0.95 | 1.57 | 1.48 | 0.88 |  |  |  |  |
| 8 | 4OAH | 1.09 | 0.76 | 0.75 | 1.44 | 1.46 | 1.07 | 0.71 |  |  |  |
| 9 | 4OAI | 1.39 | 2.09 | 1.90 | 1.92 | 1.89 | 1.37 | 1.54 | 1.87 |  |  |
| 10 | M129 | 1.66 | 1.86 | 1.78 | 1.79 | 1.80 | 1.64 | 1.85 | 1.47 | 1.65 |  |
| 11 | M133 | 1.47 | 1.73 | 1.62 | 1.92 | 1.88 | 1.44 | 1.65 | 1.22 | 0.97 | 1.14 |

**Table S4.** Mutation screening of residues on MiD51 interacting with Drp1.

| Mutations | Interactions | Mutations | Interactions |
| --- | --- | --- | --- |
| WT | / | D402R/D406R<br>/Q410T | Maintained |
| S134L/E137R/K138E/T141<br>Q | Maintained | R413E/S417Q<br>/E420R | Abolished |
| 144-RNR-146→ELE | Maintained | E441R/D444R<br>/E445R | Abolished |
| R156E/Q159T/D163R | Maintained | Y448A/Y451A | Abolished |
| E167R/S170Q/R173E | Maintained | C452S | Maintained |
| 177-PDMP-180→AAA | Maintained | E456R | Maintained |
| 182-RDMY-185→AAA | Maintained | E458R | Maintained |
| 191-YDDLQ-195→AAA | Maintained | V459E | Maintained |
| 196-VVTADH-201→AAA | Maintained | Q462F | Maintained |
| 211-EQN-213→AAA | Maintained | T463D | Maintained |
| 221-EDTIMN-226→AAA | Maintained | R234E | Weakened |
| 234-RREN-237→AAA | Weakened | E236R | Maintained |
| Δ(238-243) | Weakened | E239R | Maintained |
| Δ(234 -243) | Weakened | Y240A | Weakened |
| Y256A | Maintained | F241A | Weakened |
| K260E/D264R/K268E | Maintained | R243E | Weakened |
| 291-PPPE-294→AAA | Maintained | R413E | Maintained |
| 303-ERDK-306→AAAA | Maintained | S417Q | Maintained |
| D320R | Maintained | E420R | Abolished |
| 326-KPH-328→AAA | Maintained | E441R | Maintained |
| 329-RLAQYDN-335→AAAA | Maintained | D444R | Abolished |
| R348E/R350E/Q354T | Maintained | Y445R | Maintained |
| K372R | Maintained | Y448A | Abolished |
| 394-QEE-396→TRR | Maintained | Y451A | Abolished |

